# The Bergen Breakfast Scanning Club dataset: a deep brain imaging dataset

**DOI:** 10.1101/2023.05.30.542072

**Authors:** Meng-Yun Wang, Max Korbmacher, Rune Eikeland, Karsten Specht

## Abstract

Populational brain imaging methods based on group averages provide valuable insights into the general functions of the brain. However, they often overlook the inherent inter- and intra-subject variability, limiting our understanding of individual differences. To address this limitation, researchers have turned to big datasets and deep brain imaging datasets. Big datasets enable the exploration of inter-subject variations, while deep brain imaging datasets, involving repeated scanning of multiple subjects over time, offer detailed insights into intra-subject variability. Despite the availability of numerous big datasets, the number of deep brain imaging datasets remains limited. In this article, we present a deep brain imaging dataset derived from the Bergen Breakfast Scanning Club (BBSC) project. The dataset comprises data collected from three subjects who underwent repeated scanning over the course of approximately one year. Specifically, three types of data chunks were collected: behavioral data, functional brain data, and structural brain data. Functional brain images, encompassing magnetic resonance spectroscopy (MRS) and resting-state functional magnetic resonance imaging (fMRI), along with their anatomical reference T1-weighted brain images, were collected twice a week during the data collection period. In total, 38, 40, and 25 sessions of functional data were acquired for subjects 1, 2, and 3, respectively. On the other hand, structural brain images, including T2-weighted brain images, diffusion-weighted images (DWI), and fluid-attenuated inversion recovery (FLAIR) images, were obtained once a month. A total of 10, 9, and 6 sessions were collected for subjects 1, 2, and 3, respectively.

The primary objective of this article is to provide a comprehensive description of the data acquisition protocol employed in the BBSC project, as well as detailed insights into the preprocessing steps applied to the acquired data.

## 1. Background & Summary

One of the main goals of neuroscience is to understand the human brain’s complexity and underlying neuronal mechanisms. One common approach to achieve this goal for the last three decades was to compare two groups of people with distinct phenotypes, such as between groups (Lu et al., 2020; Lu et al., 2021; Wang, Zhang, Miao, Lin, & Yuan, 2020) or conditions (Wang, Yuan, Zhang, Xiang, & Yuan, 2020; Wang & Yuan, 2021). This group average comparison method can provide insights into the general (cross-subject) brain functional and structural organization; however, it fails to depict the intra- and inter-subject discrepancies which is essential for understanding the brain. Therefore, one needs to rethink the way how this question is approached.

Individual-level brain mapping, however, is an arising and promising measure that holds the potential to provide essential information and enhance our ability to predict behavior phenotypes. (Kong et al., 2021). In addition, it can also identify unique brain functional architecture, which is critical for the personalized medicine (Specht, 2020). There are two ways to approach individual-level brain mapping in a reliable and reproducible way: go big or go small (Gratton, Nelson, & Gordon, 2022). ‘Go big’ means big open datasets will be exploited given the consensus that very large samples (N > 1,000) are needed to reliably detect any correlation between brain measures and phenotypes (Marek et al., 2022). ‘Go small’ means several subjects will be examined multiple times to form a deep and comprehensive brain imaging dataset. While ‘go big’ can provide ample inter-subject information, ‘go small’ can delineate the in-depth intra-subject variation.

There are several big dataset initiatives (Horien et al., 2021), such as UK Biobank (Miller et al., 2016), HCP (human connectome project) (Van Essen et al., 2013), Betula (Nilsson et al., 1997), and ABIDE (autism brain imaging data exchange) (Di Martino et al., 2014), however, the initiative of deep brain imaging datasets are relatively insufficient and just getting embarking (Gratton & Braga, 2021).

Therefore, here we report a deep brain imaging dataset generated from the BBSC (Bergen Breakfast Scanning Club) project, where three subjects were repeatedly scanned for a year (Wang, Korbmacher, Eikeland, & Specht, 2022). In this project, we collected behavioral data, functional MRI (magnetic resonance imaging) data, and structural MRI data (Wang et al., 2022). It is expected that the dataset can be utilized to unveil the details of intra-subject discrepancy and can be complementary to the big datasets and contribute to the deep brain imaging dataset initiative and get the ‘go small’ approach burgeoning.

## 2. Methods

### 2.1 Participants

Three participants (**Table 1**) were scanned in this project. They are all male and can speak at least two languages (their native languages, and English). To be noticed, subject 1 has been learning another language since January 2021 and not got COVID-19 during data collection, subject 2 got COVID-19 around December 2021 while subject 3 got COVID-19 around August 2021. In addition, subject 2 did not report any long-COVID symptoms after recovery while subject 3 suffered from long-COVID after recovery mainly manifesting fatigue symptoms.

**Table 1.** Basic demographic information of subjects

|  | 1 | 2 | 3 |
| --- | --- | --- | --- |
| Gender | Male | Male | Male |
| Age | 31 | 27 | 40 |
| Laterality | Right | Right | Right |
| Regular caffeine consumption? | No | Yes | Yes |
| Regular nicotine consumption? | No | No | No |

The participants were repeatedly scanned between February 2021 and February 2022 except for the summer (Jun. to Sep. 2021) and winter (Dec. 2021 to Jan. 2022) breaks. Three protocols were implemented during data collection, which were behavioral protocol, functional protocol, and structural protocol as depicted in **Fig. 1**.

**Fig. 1.**
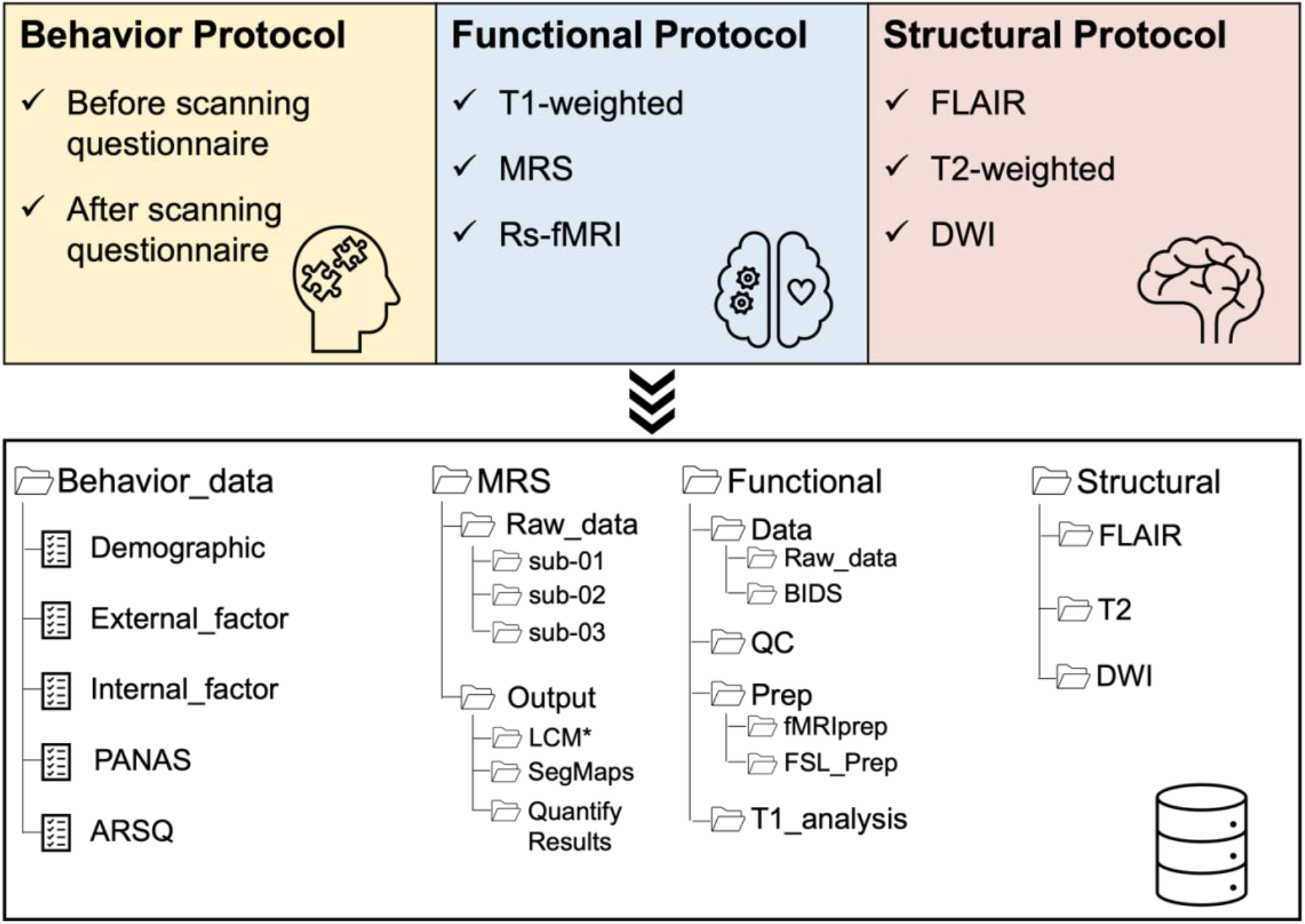
The protocols implemented in the project. The behavioral protocol was implemented along with the functional protocol with the aim of collecting exogenous and endogenous factors which could affect functional MRI data. The functional protocol was implemented aiming to collect functional MRI data including MRS data, rs-fMRI data, and their T1-weighted referenced image data. The structural protocol was implemented with the aim of collecting structural MRI data, such as T2-weighted image data, DWI, and FLAIR.

### 2.2 Behavioral Protocol

The behavior protocol collected exogenous and endogenous factors which could affect the fMRI data. Therefore, this protocol was implemented along with the functional protocol which will be described in the next section.

Before each scanning session, the participants need to fill out a questionnaire that inquiries about some basic information: the inside and outside weather information, their body statuses such as body temperature, and the expanded version of PANAS (Positive Affect and Negative Affect Schedule) scale (Watson & Clark, 1994). The PANAS-X is a 60-item scale which measures the two general affects (negative affect, positive affect) and 11 specific affects: fear, sadness, guilt, hostility, shyness, fatigue, surprise, joviality, self-Assurance, attentiveness, and serenity (Watson & Clark, 1994).

After each scanning session, the participants need to fill out another questionnaire which records information about their blood pressure and responses to the ARSQ (Amsterdam Resting-State Questionnaire) (Diaz et al., 2013). The ARSQ is a 50-item self-report survey measuring the resting-state cognition of participants, which consists of the following seven dimensions: discontinuity of mind, theory of mind, self, planning, sleepiness, comfort, and somatic awareness (Diaz et al., 2013).

The two questionnaires were digitized through the Google Forms platform, and the participants could fill them out with their mobile phones. The links to these two questionnaires are as follows: https://bit.ly/3DcVurg; https://bit.ly/3R6fMJ0.

### 2.3 Functional Protocol

This protocol aims to collect functional MRI data including MRS (magnetic resonance spectroscopy) data, (resting state) rs-fMRI data, and their anatomical reference T1-weighted (T1w) MRI data, which lasts around 25 mins in total. MRI data were collected with a 3T MR scanner (GE Discovery MR750) with a 32-channel head coil at the Haukeland University Hospital in Bergen, Norway. The protocol was implemented once or twice a week during the data collection. The technical details are as follows.

First, around seven-minute (7min and 7s) structural T1w image data were acquired using a 3D FSPGR (Fast Spoiled Gradient-Recalled Echo) sequence with the following parameters: 188 contiguous slices acquired, with TR (repetition time) = 6.88 ms, TE (echo time) = 2.95 ms, FA (flip angle) = 12°, slice thickness = 1 mm, in-plane resolution = 1 mm × 1 mm, and FOV (field of view) = 256 mm, with an isotropic voxel size of 1 mm^3^.

Second, around four-minute (3 min and 48s) ^1^H-MRS-spectra data were obtained from the anterior cingulate cortex (ACC) (voxel size 25 × 25 × 25 mm) by using a single-voxel point-resolved spectroscopy (PRESS) sequence (TE/TR = 35 ms/1,500 ms, 128 repetitions). Unsuppressed water reference spectra (eight repetitions) were acquired automatically after the water-suppressed metabolite spectra.

Last, around twelve-minute (12 mins and 10s) rs-fMRI data containing a total of 350 brain volumes were collected using gradient EPI (echo-planar imaging) pulse sequence with the following parameters: 38 slices with a 0.5 mm gap between slices acquired from bottom to top, TR = 2.05 s, TE = 30 ms, FA = 90°, slice thickness = 3.5 mm, in-plane resolution = 1.72 mm × 1.72 mm, FOV = 220 mm, with a voxel size of 1.72 mm × 1.72 mm × 3 mm. The respiratory and pulse data for each session were also collected along with the rs-fMRI data. During the data acquisition, the participants were instructed to lie in the scanner still and relaxed with their eyes closed but without falling asleep.

The MRS raw data were in *MRS/Raw_data* while the T1w and resting state brain image data were in the *Functional/Data/Raw_data* folder (**Fig. 1**).

### 2.4 Structural Protocol

This protocol aims to collect structural MRI data including T2-weighted MRI data, DWI (diffusion-weighted image) data, and FLAIR (Fluid attenuated inversion recovery) data which lasts around 25 mins in total. MRI data were collected with the same MR scanner (GE Discovery MR750) with a 32-channel head coil at the Haukeland University Hospital in Bergen, Norway. The protocol was implemented once a month during data collection. The data collected from the protocol were organized in the *Structure* folders and the technical details of data collection are as follows.

Around seven-minute (6 mins and 28s) FLAIR image data were first acquired using a 3D Sag Cube T2 FLAIR sequence with the following parameters: 360 contiguous slices acquired, with TR = 8002 ms, TE = 97 ms, FA = 90°, slice thickness = 0.5 mm, in-plane resolution = 0.5 mm × 0.5 mm, and FOV = 256 mm.

Around five-minute (4 mins and 28s) T2-weighted image data were then acquired using a 3D Sag T2 Cube sequence with the following parameters: 174 contiguous slices acquired, with TR = 3002 ms, TE = 109 ms, FA = 90°, slice thickness = 1 mm, in-plane resolution = 0.5 mm × 0.5 mm, and FOV = 256 mm.

Around ten-minute DWI data were acquired using a 2D Spin Echo sequence with the following parameters: b2800 (5mins) and b1000 (5mins); 67 contiguous slices acquired, with TR = 8000 ms, TE = 79.4 ms, FA = 90°, slice thickness = 2 mm, in-plane resolution = 1 mm × 1 mm, and FOV = 256 mm.

## 3. Data Records (*Preprocessed Data*)

### 3.1 Curated Data from Behavior protocol

The curated behavior data were in the *Behavior_data* folder compassing the following five files (**Fig. 1**).

The *Demographic* file describes the basic information about the subjects, such as age, sex, ethnicity, et al.

The *External_factor* file records the outside factors before, during, or after the scanning, such as the outside temperature and humidity, the scanner room temperature and humidity, the control room temperature and humidity, daytime length, scanning timestamp (morning or afternoon or evening; in which month), et al.

The *Internal_factor* file stores the endogenous factors before, during, or after scanning, such as the body temperature, blood pressure before and after scanning, drowsiness during scanning, length of sleep the day before the scanning, slept or not during scanning, et al.

The *PANAS* file contains the responses measured with the PANAS-X (Watson & Clark, 1994) before scanning.

The *ARSQ* file records the participants’ responses measured with the Amsterdam Resting-State Questionnaire (Diaz et al., 2013) after the scanning.

### 3.2 Pre/processed Data from Functional protocol

In total, 38, 40, and 25 sessions were collected for subjects 1, 2, and 3, respectively. The data collected from the functional protocol were organized into two directories: the *MRS* and *Functional* folders (**Fig. 1**).

The *MRS* folder contains all the raw and analyzed MRS data while the *Functional* folder includes all the rs-fMRI and T1w data compassing four subfolders: *Data, QC, T1_analysis*, and *Prep*. The *Data* subfolder includes two directories: the raw data (*Raw_data*) in DICOM (digital imaging and communications in medicine) format and the *BIDS* data in NIfTI (neuroimaging informatics technology initiative) format following the BIDS (brain imaging data structure) structure (Gorgolewski et al., 2016). The *QC* subfolder includes all the output matrices of the quality check using MRIQC 16.0.1 (Esteban et al., 2017). The *Prep* subfolder includes two directories: the data (*fMRIprep*) preprocessed with fMRIPrep 20.0.1 (Esteban et al., 2019) and the data (*FSL_Prep*) preprocessed with FSL 6.0.5.2:dc6f4207 (Jenkinson, Bannister, Brady, & Smith, 2002). The *T1_analysis* subfolder contains all the output of the T1w brain image processed with FreeSurfer 7.2.0 (Dale, Fischl, & Sereno, 1999; Fischl, Sereno, & Dale, 1999). The details of data preprocessing are described as follows.

#### 3.2.1 MRS

The MRS data were analyzed with Osprey 1.1.0 (Oeltzschner et al., 2020) with the integrated LCModel fitting algorithm (Provencher, 1993) based on the MATLAB platform, which provides an automated and uniform processing pipeline including pre-processing, linear combination modeling, tissue correction, and quantification. The processed MRS data were in the *MRS/output* folder.

A simple description of the processing pipeline is as follows. The raw data were first aligned and averaged (**Fig. 2A**), and then fitted using the LCModel, which is embedded in the Osprey, with the default settings across a frequency range of 0.5 to 4.0 ppm (parts per million). 22 metabolites and 9 macromolecular/lipids were included in the model: Asc (ascorbate), Asp (aspartate), Cr (creatine), GABA (gamma-Aminobutyric acid), GPC (glycerophosphocholine), GSH (glutathione), Gln (glutamine), Glu (glutamate), Ins, Lac (lactose), NAA (N-acetylaspartate), NAAG (N-acetyl-aspartyl-glutamate), PCh (phosphocholine), PCr (phosphocreatine), PE (Phosphorylethanolamine), SCyllo (scyllo-inositol), Tau, CrCH2 (**Fig. 2B**). Before quantification, the brain was segmented into GM (grey matter), WM (white matter), and CSF (cerebrospinal fluid) after coregistration to the structural image with functional from SPM12 (Friston et al., 1994) emulated by Osprey. Quantification of the metabolites was calculated in four different ways: as the ratio to tCr (total creatine); as water-scaled metabolite estimates; as water-scaled metabolite estimates corrected for the volume fraction of CSF; and as fully tissue-and-relaxation-corrected molal concentration estimates. For more details, we recommend interested readers to the original article about Osprey (Oeltzschner et al., 2020).

**Fig. 2.**
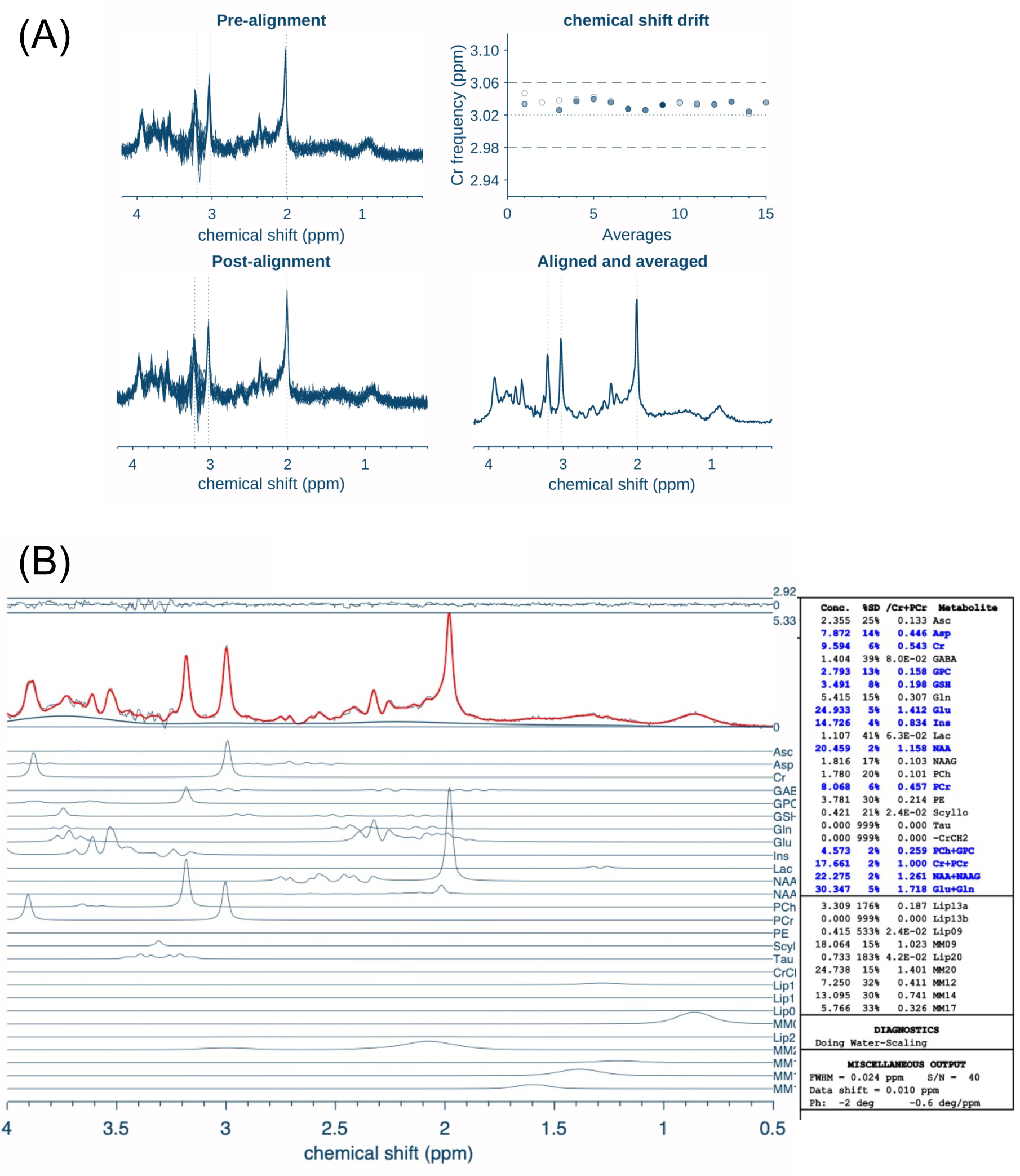
Preprocessing of MRS with Osprey. (A). The raw data were aligned and averaged. (B) Then the data were fitted using LCModel embedded in Osprey. The metabolites in blue color mean the estimation is reliable (criteria: %SD <15%). Figures were generated with Osprey.

The modeling results were in the *MRS/output/LCM\** folder; the segmentation results were in the *MRS/output/SegMaps* folder; the quantification results were in the *MRS/output/QuantifyResults* folder (**Fig. 1**).

#### 3.2.2 rs-fMRI data

The raw DICOM data were converted into NIfTI format with *dcm2niix* (v1.0.20211006) (Li, Morgan, Ashburner, Smith, & Rorden, 2016) and reorganized into BIDS (Gorgolewski et al., 2016) structure which were in the *Functional/Data/BIDS* folder. The data were then quality checked with MRIQC 0.16.1 (Esteban et al., 2017) and the outputs of the quality check were in the *Functional/QC* folder (**Fig. 1**). The details of the quality check can be found in the Technical Validation part. The data were then preprocessed with two preprocessing pipelines, one with fMRIPrp and one with FEAT (FMRI Expert Analysis Tool) embedded in the FSL (FMRIB’s Software Library, www.fmrib.ox.ac.uk/fsl) (Smith et al., 2004).

##### Preprocessed with fMRIPrep

The data were preprocessed with *fMRIPrep* 22.0.1 (Esteban et al., 2019), which is based on *Nipype* 1.8.4 (Gorgolewski et al., 2011).

###### Anatomical data preprocessing

A total of 38, 40, and 25 T1w images for subjects 1, 2, and 3 were preprocessed. All of them were first corrected for intensity non-uniformity (Tustison et al., 2010) and a T1w-reference map was computed after the registration of all (within-subject) T1w images (Dale et al., 1999; Fischl et al., 1999; Reuter, Rosas, & Fischl, 2010). The T1w-reference was then skull-stripped and brain tissue segmentation of CSF, WM, and GM was performed on the brain-extracted T1w (Zhang, Brady, & Smith, 2001). Volume-based spatial normalization to two standard spaces (MNI152NLin6Asym, MNI152NLin2009cAsym) was performed through nonlinear registration using brain-extracted versions of both the T1w reference and the T1w template.

###### Functional data preprocessing

The following preprocessing steps were performed for each of the BOLD runs obtained from each subject. First, a reference volume and its skull-stripped version were generated and head-motion parameters were estimated before any spatiotemporal filtering (Jenkinson et al., 2002; Smith et al., 2004). BOLD runs were slice-time corrected to 1s (Cox & Hyde, 1997) and were resampled onto their original, native space by applying the transforms to correct for head-motion, which was the preprocessed BOLD in native space.

The BOLD reference was then co-registered to the T1w reference with the boundary-based registration (Greve & Fischl, 2009) with six degrees of freedom. Several confounding time-series were calculated based on the preprocessed BOLD in native space: FD (framewise displacement), DVARS (D referring to a temporal derivative of time courses, VARS referring to RMS variance over voxels) (Power, Barnes, Snyder, Schlaggar, & Petersen, 2012; Power et al., 2014), and three region-wise global signals (CSF, WM, and the whole brain).

Additionally, a set of physiological regressors were extracted to allow for component-based noise corrections (CompCor) (Behzadi, Restom, Liau, & Liu, 2007). The calculated FD, DVARS, global signals, and ComCor values were placed within the corresponding confounds file. The BOLD time-series were resampled into standard space, generating a preprocessed BOLD run in MNI152NLin6Asym space.

The details of the fMRIPrep preprocessing are described in the **supplementary materials** and the final preprocessed data are illustrated in **Fig. 3**. The preprocessed data were *BBSC/Functional/Prep/fMRIprep/sub-ID/ses-ID/ sub-ID_ses-ID_task-rest_space-MNI152NLin6Asym_res-2_desc-preproc_bold.nii.gz*

**Fig. 3.**
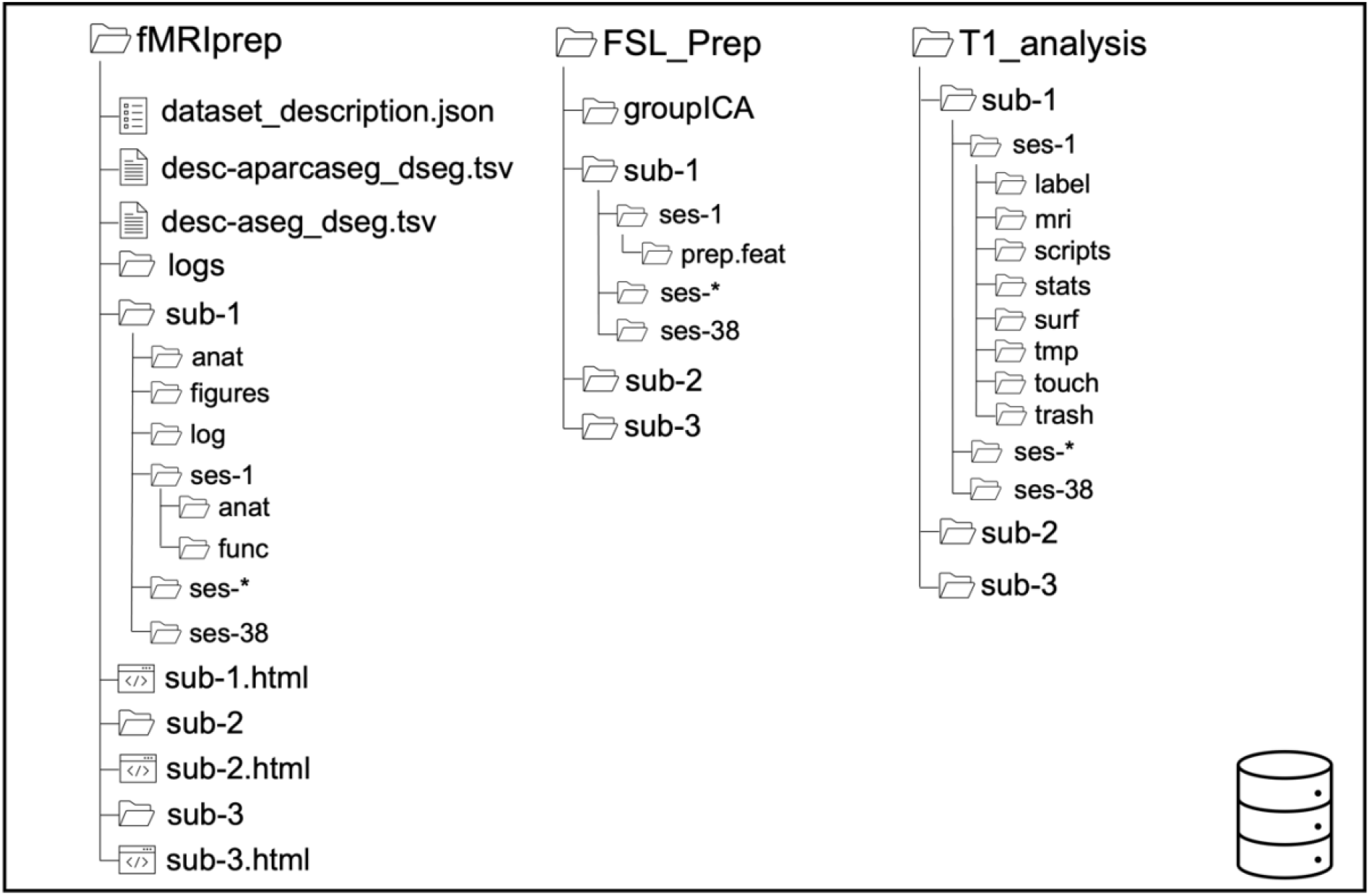
The data structure of the preprocessed data.

##### Preprocessed with FSL

Besides the rs-fMRI data were preprocessed with fMRIPrep, they were also preprocessed by the FEAT (Version 6.00) with default parameters, which is a part of the FSL with version 6.0.5.2:dc6f4207 (Jenkinson et al., 2002).

The following processing was applied: registration to high-resolution structural and/or standard space images was carried out using FLIRT (Jenkinson et al., 2002; Jenkinson & Smith, 2001); motion correction using MCFLIRT (Jenkinson et al., 2002); slice-timing correction using Fourier-space time-series phase-shifting; non-brain removal using BET (Smith, 2002); spatial smoothing using a Gaussian kernel of FWHM 5mm; grand-mean intensity normalization of the entire 4D dataset by a single multiplicative factor; high-pass temporal filtering (Gaussian-weighted least-squares straight line fitting, with sigma = 50.0s). ICA-based exploratory data analysis was carried out using MELODIC (Beckmann & Smith, 2004), in order to investigate the possible presence of unexpected artifacts or activation. The preprocessed data in native space were *BBSC/Functional/Prep/FSL_Prep/sub-ID/ses-ID/prep.feat/ filtered_func_data.nii.gz*. Then the data were cleaned with FIX (“FMRIB’s ICA-based X-noiseifier”) (Salimi-Khorshidi et al., 2014), part of FSL, and were standardized to the MNI152NLin6Asym template with the FSL function *applywarp*. The preprocessed data were in the FSL_Prep folder as illustrated in **Fig. 3**. The cleaned data in standard space (MNI152NLin6Asym) were *BBSC/Functional/Prep/FSL_Prep/sub-ID/ses-ID/prep.feat/ filtered_func_data_clean_standard.nii.gz*

#### 3.2.3 T1-weighted image

The T1w images were preprocessed with the command *recon-all* by using FreeSurfer (version: freesurfer-darwin-macOS-7.2.0-20210713-aa8f76b) (Dale et al., 1999; Fischl et al., 1999) with default parameters. The preprocessing and the pipeline are described somewhere else in the official document (https://surfer.nmr.mgh.harvard.edu/fswiki/FreeSurferAnalysisPipelineOverview). Briefly speaking, the preprocessing includes skull-stripping, white mattering segmentation, surface extraction, labeling, and surface atlas (such as fsaverage) registration. The results were in the *Functional/T1_analysis* folder (**Fig. 3**).

### 3.3 Preprocessed Data from Structural protocol

In total, 10, 9, and 6 sessions were collected for subjects 1, 2, and 3, respectively. The data collected from the structural protocol were in the *Structural* folder.

The structural data will be reported separately.

## 4. Technical Validation

### MRS data

MRS data have already been quality checked with Osprey described in the previous sections. Those metabolisms in blue color (**Fig. 2B**) have passed the quality check (%SD < 15%) and can be used in further analysis, which are Asp, Cr, GPC, GSH, Glu, Ins, NAA, PCr, tCho (Pch+Gpc), tCr (Cr+PCr), tNAA (NAA+NAAG), and Glx (Glu+Gln).

The rs-fMRI and T1w data were quality checked with MRIQC 0.16.1 (Esteban et al., 2017) and naked eyes, and the outputs of the quality check were in the *Functional/QC* folder (**Fig. 1**). The pipeline is illustrated in **Figs. 4-5**.

**Fig. 4.**
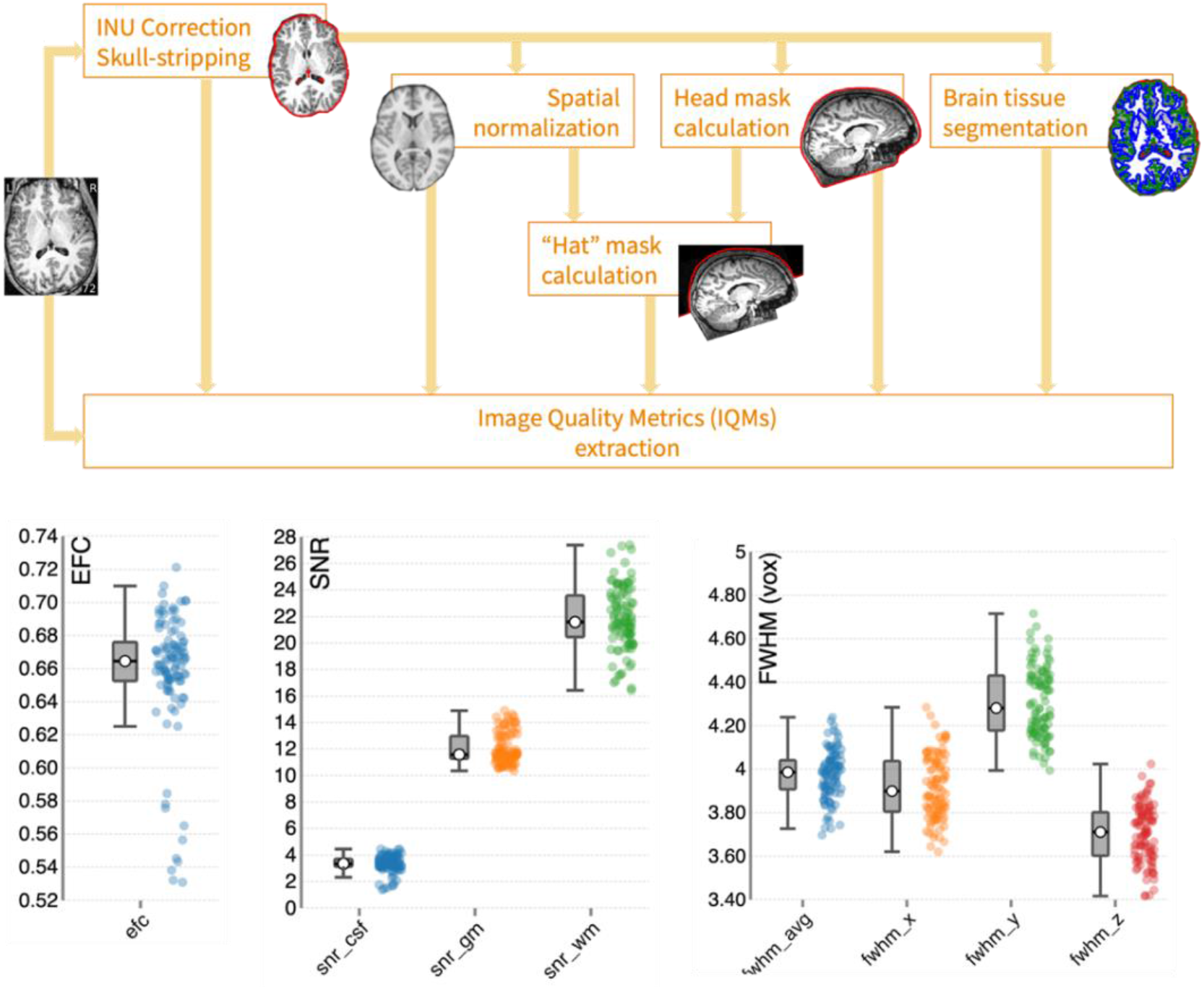
The pipeline and outputs of the MRIQC in checking T1w images. The upper chart is the pipeline while the bottom charts are the distributions of the data quality indices of the BBSC dataset. The average EFC and FWHM values are 0.67 and 4, of which the lower the better. The average SNR (grey matter) value is 12, of which the higher the better. Compared with other big datasets, such as ABIDE (EFC = 0.43; FWHM = 5.73; SNR = 16), the BBSC dataset manifests good data quality. Figures were generated with the MRIQC software (Esteban et al., 2017).

### T1w data with good data quality

Regarding the T1w data, the values of the following three parameters were used to indicate the image quality (**Fig. 4**.). The FWHM (full-width half maximum) measures the FWHM of the spatial distribution of the image intensity values in voxel units representing the smoothness of voxels, of which the lower values are better (Friedman, Glover, Krenz, Magnotta, & First, 2006). The EFC (Entropy Focus Criterion) calculates the Shannon entropy of voxel intensities proportional to the maximum possible entropy for a similarly sized image indicating ghosting and head motion-induced blurring (Atkinson, Hill, Stoyle, Summers, & Keevil, 1997), of which the lower values are better. The SNR (Signal-to-Noise Ratio) measures the mean intensity within brain tissues divided by the standard deviation of the values outside the brain, of which the higher values are better (Magnotta, Friedman, & First, 2006).

The FWHM and EFC values reflect the technical quality such as the MRI system and parameters while the SNR values are reflections of the physiological factors such as the head motions (Shehzad et al., 2015). As illustrated in **Fig. 4**, the average EFC and FWHM values of the BBSC dataset are 0.67 and 4, respectively, while the average SNR value of the grey matter is around 12, which are comparable to other big datasets, such as the ABIDE data set (EFC = 0.43, FWHM = 5.73, and SNR = 16) (Shehzad et al., 2015). The distributions of the three parameters are clustered without any significant outliers, which indicates that the data quality of the T1w brain imaging is good.

### rs-fMRI with good motion control

Regarding the rs-fMRI data, the following parameters were considered to measure the data quality (**Fig. 5**). The standardized DVARS (D referring to a temporal derivative of time courses, VARS referring to RMS variance over voxels) measures the average change in mean intensity in comparison to the previous timepoint, with which the lower values are better (Power et al., 2012). The FD (framewise displacement) compares the head motions between the current and previous volumes, which is calculated by summing the absolute value of displacement changes in the x y, and z directions and rotational changes about those three axes. The rotational changes are given distance values based on the changes across the surface of an 80 mm radius sphere. The lower values of FD the better (Power et al., 2012).

**Fig. 5.**
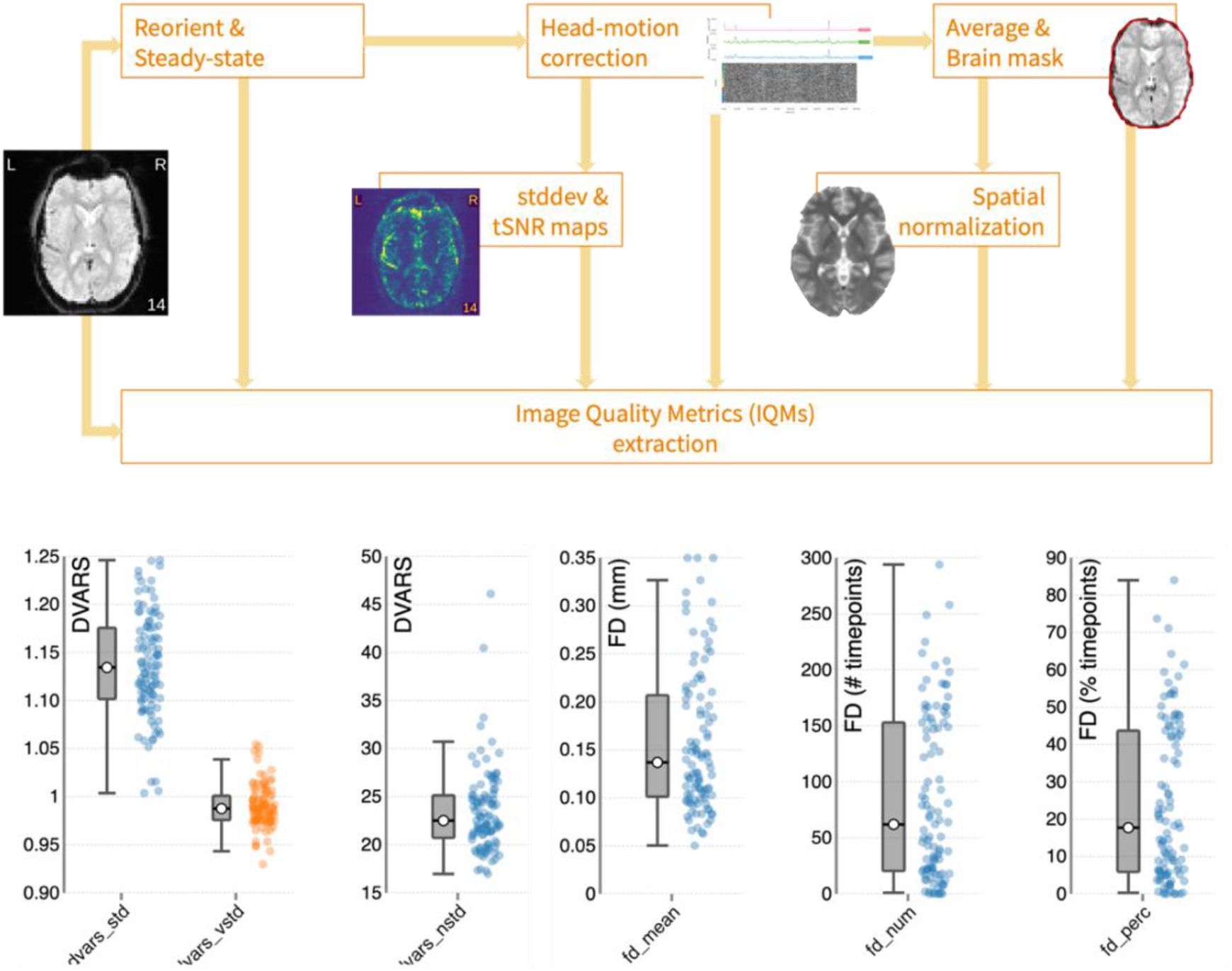
The pipeline and outputs of the MRIQC in checking rs-fMRI data. The upper chart is the pipeline while the bottom charts are the distributions of the data quality indices of the BBSC dataset. The average DVARS value is 1.14 (the lower the better). The average FD value is 0.14 (the lower the better). The number and percentage of volumes exceeding 0.2mm are shown, in which only around 15 sessions are over 50 %. The detailed information on each subject can be found in the QC folder. Figures were generated with MRIQC software (Esteban et al., 2017).

The average DVARS value is 1.14 (the lower the better). The average FD value is 0.14 mm (the lower the better). The number and percentage of volumes exceeding 0.2 mm are shown in **Fig. 5**, in which only around 15 out of 103 sessions are over 50 %. Those results show that the rs-fMRI data were in good control of head motions.

### Usage Note

The internal_factor file records whether the participants fall asleep during the data collection or not, please pay attention, where 1 represents falling asleep, 0 represents awake, and 0.5 represents maybe falling asleep (the participants were not 100% sure).

Regarding the MRS data, the first session of subject 2 did not record any MRS data. So, when correlating MRS data to the fMRI data in subject 2, please remember the first record of MRS data should be corresponding to the second session, and so on and so forth.

Regarding the rs-fMRI data, the sessions exceeding the FD=0.2 mm with 50% are mostly clustered in subject 3.

## Supporting information

The details of the fMRIPrep preprocessing are described in the supplementary materials

## Data and code availability

All the data were uploaded into the OpenNeuro platform (LINK). All the code is accessible on the GitHub platform (LINK).

## Funding

This study was financed by the Research Council of Norway (Project number: 276044: When default is not default: Solutions to the replication crisis and beyond).

## Acknowledgments

We thank Alexandra Vik and Liucija Vaisvilaite for their suggestions on the post resting-state fMRI data collection questionnaire. We also thank Alexander R. Craven for his expertise in MRS data analysis. Besides, we thank Ulvhild Faerøvik and Tania Martínez-Montero for providing useful resources. More importantly, we appreciate the technical support and data collection support from our radiologists (Christel Jansen, Eva Øksnes, Roger Barndon, Trond Øvreaas, Tor Fjørtoft, and Turid Randa) at the Haukeland University Hospital.

## Notes

### Competing Interest Statement

The authors have declared no competing interest.

### Summary of Updates

The abstract part is updated.

## References

Atkinson, D., Hill, D. L., Stoyle, P. N., Summers, P. E., & Keevil, S. F. (1997). Automatic correction of motion artifacts in magnetic resonance images using an entropy focus criterion. Ieee Transactions on Medical Imaging, 16(6), 903–910. doi:10.1109/42.650886

Beckmann, C. F., & Smith, S. A. (2004). Probabilistic independent component analysis for functional magnetic resonance imaging. Ieee Transactions on Medical Imaging, 23(2), 137–152. doi:10.1109/Tmi.2003.822821

Behzadi, Y., Restom, K., Liau, J., & Liu, T. T. (2007). A component based noise correction method (CompCor) for BOLD and perfusion based fMRI. Neuroimage, 37(1), 90–101. doi:10.1016/j.neuroimage.2007.04.042

Cox, R. W., & Hyde, J. S. (1997). Software tools for analysis and visualization of fMRI data. Nmr in Biomedicine, 10(4-5), 171–178. doi:Doi 10.1002/(Sici)1099-1492(199706/08)10:4/5<171::Aid-Nbm453>3.0.Co;2-L

Dale, A. M., Fischl, B., & Sereno, M. I. (1999). Cortical surface-based analysis. I. Segmentation and surface reconstruction. Neuroimage, 9(2), 179–194. doi:10.1006/nimg.1998.0395

Di Martino, A., Yan, C. G., Li, Q., Denio, E., Castellanos, F. X., Alaerts, K., … Milham, M. P. (2014). The autism brain imaging data exchange: towards a large-scale evaluation of the intrinsic brain architecture in autism. Molecular Psychiatry, 19(6), 659–667. doi:10.1038/mp.2013.78

Diaz, B. A., Van Der Sluis, S., Moens, S., Benjamins, J. S., Migliorati, F., Stoffers, D., … Linkenkaer-Hansen, K. (2013). The Amsterdam Resting-State Questionnaire reveals multiple phenotypes of resting-state cognition. Frontiers in Human Neuroscience,7, 446. doi:10.3389/fnhum.2013.00446

Esteban, O., Birman, D., Schaer, M., Koyejo, O. O., Poldrack, R. A., & Gorgolewski, K. J. (2017). MRIQC: Advancing the automatic prediction of image quality in MRI from unseen sites. PLoS One, 12(9), e0184661. doi:10.1371/journal.pone.0184661

Esteban, O., Markiewicz, C. J., Blair, R. W., Moodie, C. A., Isik, A. I., Erramuzpe, A., … Gorgolewski, K. J. (2019). fMRIPrep: a robust preprocessing pipeline for functional MRI. Nature Methods, 16(1), 111–116. doi:10.1038/s41592-018-0235-4

Fischl, B., Sereno, M. I., & Dale, A. M. (1999). Cortical surface-based analysis. II: Inflation, flattening, and a surface-based coordinate system. Neuroimage, 9(2), 195–207. doi:10.1006/nimg.1998.0396

Friedman, L., Glover, G. H., Krenz, D., Magnotta, V., & First, B. (2006). Reducing inter-scanner variability of activation in a multicenter fMRI study: role of smoothness equalization. Neuroimage, 32(4), 1656–1668. doi:10.1016/j.neuroimage.2006.03.062

Friston, K. J., Holmes, A. P., Worsley, K. J., Poline, J. P., Frith, C. D., & Frackowiak, R. S. (1994). Statistical parametric maps in functional imaging: a general linear approach. Human Brain Mapping, 2(4), 189–210.

Gorgolewski, K. J., Auer, T., Calhoun, V. D., Craddock, R. C., Das, S., Duff, E. P., … Poldrack, R. A. (2016). The brain imaging data structure, a format for organizing and describing outputs of neuroimaging experiments. Scientific Data, 3. doi:ARTN 160044 10.1038/sdata.2016.44

Gorgolewski, K. J., Burns, C. D., Madison, C., Clark, D., Halchenko, Y. O., Waskom, M. L., & Ghosh, S. S. (2011). Nipype: a flexible, lightweight and extensible neuroimaging data processing framework in python. Front Neuroinform,5, 13. doi:10.3389/fninf.2011.00013

Gratton, C., & Braga, R. M. (2021). Editorial overview: Deep imaging of the individual brain: past, practice, and promise. Current Opinion in Behavioral Sciences,40, Iii–Vi. doi:10.1016/j.cobeha.2021.06.011

Greve, D. N., & Fischl, B. (2009). Accurate and robust brain image alignment using boundary-based registration. Neuroimage, 48(1), 63–72. doi:10.1016/j.neuroimage.2009.06.060

Horien, C., Noble, S., Greene, A. S., Lee, K., Barron, D. S., Gao, S. Y., … Scheinost, D. (2021). A hitchhiker’s guide to working with large, open-source neuroimaging datasets. Nature Human Behaviour, 5(2), 185–193. doi:10.1038/s41562-020-01005-4

Jenkinson, M., Bannister, P., Brady, M., & Smith, S. (2002). Improved optimization for the robust and accurate linear registration and motion correction of brain images. Neuroimage, 17(2), 825–841. doi:10.1016/s1053-8119(02)91132-8

Jenkinson, M., & Smith, S. (2001). A global optimisation method for robust affine registration of brain images. Medical Image Analysis, 5(2), 143–156. doi:Doi 10.1016/S1361-8415(01)00036-6

Kong, R., Yang, Q., Gordon, E., Xue, A. H., Yan, X. X., Orban, C., … Yeo, B. T. T. (2021). Individual-Specific Areal-Level Parcellations Improve Functional Connectivity Prediction of Behavior. Cerebral Cortex, 31(10), 4477–4500. doi:10.1093/cercor/bhab101

Li, X., Morgan, P. S., Ashburner, J., Smith, J., & Rorden, C. (2016). The first step for neuroimaging data analysis: DICOM to NIfTI conversion. Journal of Neuroscience Methods,264, 47–56. doi:10.1016/j.jneumeth.2016.03.001

Lu, F. M., Liu, P. Q., Chen, H., Wang, M. Y., Xu, S. Y., Yuan, Z., … Zhou, J. S. (2020). More than just statics: Abnormal dynamic amplitude of low-frequency fluctuation in adolescent patients with pure conduct disorder. Journal of Psychiatric Research,131, 60–68. doi:10.1016/j.jpsychires.2020.08.027

Lu, F. M., Wang, M. Y., Xu, S. Y., Chen, H., Yuan, Z., Luo, L. Z., … Zhou, J. S. (2021). Decreased interhemispheric resting-state functional connectivity in male adolescents with conduct disorder. Brain Imaging and Behavior, 15(3), 1201–1210. doi:10.1007/s11682-020-00320-8

Magnotta, V. A., Friedman, L., & First, B. (2006). Measurement of Signal-to-Noise and Contrast-to-Noise in the fBIRN Multicenter Imaging Study. Journal of Digital Imaging, 19(2), 140–147. doi:10.1007/s10278-006-0264-x

Miller, K. L., Alfaro-Almagro, F., Bangerter, N. K., Thomas, D. L., Yacoub, E., Xu, J. Q., … Smith, S. M. (2016). Multimodal population brain imaging in the UK Biobank prospective epidemiological study. Nature Neuroscience, 19(11), 1523–1536. doi:10.1038/nn.4393

Nilsson, L. G., Backman, L., Erngrund, K., Nyberg, L., Adolfsson, R., Bucht, G., … Winblad, B. (1997). The Betula prospective cohort study: Memory, health and aging. Aging Neuropsychology and Cognition, 4(1), 1–32. doi:Doi 10.1080/13825589708256633

Oeltzschner, G., Zollner, H. J., Hui, S. C. N., Mikkelsen, M., Saleh, M. G., Tapper, S., & Edden, R. A. E. (2020). Osprey: Open-source processing, reconstruction & estimation of magnetic resonance spectroscopy data. Journal of Neuroscience Methods, 343. doi:10.1016/j.jneumeth.2020.108827

Power, J. D., Barnes, K. A., Snyder, A. Z., Schlaggar, B. L., & Petersen, S. E. (2012). Spurious but systematic correlations in functional connectivity MRI networks arise from subject motion. Neuroimage, 59(3), 2142–2154. doi:10.1016/j.neuroimage.2011.10.018

Power, J. D., Mitra, A., Laumann, T. O., Snyder, A. Z., Schlaggar, B. L., & Petersen, S. E. (2014). Methods to detect, characterize, and remove motion artifact in resting state fMRI. Neuroimage,84, 320–341. doi:10.1016/j.neuroimage.2013.08.048

Provencher, S. W. (1993). Estimation of metabolite concentrations from localized in vivo proton NMR spectra. Magnetic Resonance in Medicine, 30(6), 672–679. doi:10.1002/mrm.1910300604

Reuter, M., Rosas, H. D., & Fischl, B. (2010). Highly accurate inverse consistent registration: a robust approach. Neuroimage, 53(4), 1181–1196. doi:10.1016/j.neuroimage.2010.07.020

Salimi-Khorshidi, G., Douaud, G., Beckmann, C. F., Glasser, M. F., Griffanti, L., & Smith, S. M. (2014). Automatic denoising of functional MRI data: combining independent component analysis and hierarchical fusion of classifiers. Neuroimage,90, 449–468. doi:10.1016/j.neuroimage.2013.11.046

Shehzad, Z., Giavasis, S., Li, Q., Benhajali, Y., Yan, C., Yang, Z., … Craddock, C. (2015). The Preprocessed Connectomes Project Quality Assessment Protocol-a resource for measuring the quality of MRI data. Frontiers in neuroscience, 47.

Smith, S. M. (2002). Fast robust automated brain extraction. Human Brain Mapping, 17(3), 143–155. doi:10.1002/hbm.10062

Smith, S. M., Jenkinson, M., Woolrich, M. W., Beckmann, C. F., Behrens, T. E. J., Johansen-Berg, H., … Matthews, P. M. (2004). Advances in functional and structural MR image analysis and implementation as FSL. Neuroimage,23, S208–S219. doi:10.1016/j.neuroimage.2004.07.051

Specht, K. (2020). Current Challenges in Translational and Clinical fMRI and Future Directions. Frontiers in Psychiatry, 10. doi:10.3389/fpsyt.2019.00924

Tustison, N. J., Avants, B. B., Cook, P. A., Zheng, Y. J., Egan, A., Yushkevich, P. A., & Gee, J. C. (2010). N4ITK: Improved N3 Bias Correction. Ieee Transactions on Medical Imaging, 29(6), 1310–1320. doi:10.1109/Tmi.2010.2046908

Van Essen, D. C., Smith, S. M., Barch, D. M., Behrens, T. E. J., Yacoub, E., Ugurbil, K., & Consortium, W.-M. H. (2013). The WU-Minn Human Connectome Project: An overview. Neuroimage,80, 62–79. doi:10.1016/j.neuroimage.2013.05.041

Wang, M. Y., Korbmacher, M., Eikeland, R., & Specht, K. (2022). Deep brain imaging of three participants across 1 year: The Bergen breakfast scanning club project. Frontiers in Human Neuroscience, 16. doi:ARTN 1021503 10.3389/fnhum.2022.1021503

Wang, M. Y., Yuan, A. Z., Zhang, J., Xiang, Y. T., & Yuan, Z. (2020). Functional near-infrared spectroscopy can detect low-frequency hemodynamic oscillations in the prefrontal cortex during steady-state visual evoked potential-inducing periodic facial expression stimuli presentation. Visual Computing for Industry Biomedicine and Art, 3(1). doi:ARTN 28 10.1186/s42492-020-00065-7

Wang, M. Y., & Yuan, Z. (2021). EEG Decoding of Dynamic Facial Expressions of Emotion: Evidence from SSVEP and Causal Cortical Network Dynamics. Neuroscience,459, 50–58. doi:10.1016/j.neuroscience.2021.01.040

Wang, M. Y., Zhang, Z. M., Miao, X. C., Lin, X. H., & Yuan, Z. (2020). Electrophysiological Evidence of Attentional Avoidance in Sub-Clinical Individuals With Obsessive-Compulsive Symptoms. Ieee Access,8, 91020–91027. doi:10.1109/Access.2020.2994452

Watson, D., & Clark, L. A. (1994). The PANAS-X: Manual for the positive and negative affect schedule-expanded form. doi:10.17077/48vt-m4t2

Zhang, Y. Y., Brady, M., & Smith, S. (2001). Segmentation of brain MR images through a hidden Markov random field model and the expectation-maximization algorithm. Ieee Transactions on Medical Imaging, 20(1), 45–57. doi:Doi 10.1109/42.906424

